# Infectious Entry of Merkel Cell Polyomavirus

**DOI:** 10.1101/456673

**Authors:** Miriam Becker, Melissa Dominguez, Lilo Greune, Laura Soria-Martinez, Moritz M. Pfleiderer, Rachel Schowalter, Christopher B. Buck, Bärbel S. Blaum, M. Alexander Schmidt, Mario Schelhaas

**Affiliations:** Institute of Cellular Virology, ZMBE, University of Münster, Germany; Cluster of Excellence EXC1003, Cells in Motion, CiM, Münster, Germany; Institute of Infectiology, ZMBE, University of Münster, Germany; Research Group, FOR2327 “ViroCarb”, Coordinating University of Tübingen, Germany; Interfaculty Institute of Biochemistry (IFIB), University of Tübingen, Germany; Center for Cancer Research, National Cancer Institute, Bethesda, MD, USA

**Keywords:** polyomavirus, MCPyV, virus entry, virus-host interaction, endocytosis

## Abstract

Merkel Cell Polyomavirus (MCPyV) is a small, non-enveloped tumor virus associated with an aggressive form of skin cancer, the Merkel cell carcinoma (MCC). MCPyV infections are highly prevalent in the human population with MCPyV virions being continuously shed from human skin. However, the precise host cell tropism(s) of MCPyV remains unclear: MCPyV is able to replicate within a subset of dermal fibroblasts, but MCPyV DNA has also been detected in a variety of other tissues. However, MCPyV appears different from other polyomaviruses as it requires sulfated polysaccharides such as heparan sulfates and/or chondroitin sulfates for initial attachment. Like other polyomaviruses, MCPyV engages sialic acid as a (co-receptor). To explore the infectious entry process of MCPyV, we analyzed the cell biological determinants of MCPyV entry into A549 cells, a highly transducible lung carcinoma cell line, in comparison to well-studied simian virus 40 and a number of other viruses. Our results indicate that MCPyV enters cells via caveolar/lipid raft-mediated endocytosis but not macropinocytosis, clathrin-mediated endocytosis or glycosphingolipid-enriched carriers. The viruses internalized in small endocytic pits that led the virus to endosomes and from there to the endoplasmic reticulum (ER). Similar to other polyomaviruses, trafficking required microtubular transport, acidification of endosomes, and a functional redox environment. To our surprise, the virus was found to acquire a membrane envelope within endosomes, a phenomenon not reported for other viruses. Only minor amounts of viruses reached the ER, while the majority was retained in endosomal compartments suggesting that endosome-to-ER trafficking is a bottleneck during infectious entry.

**Importance:** MCPyV is the first polyomavirus directly implicated in the development of an aggressive human cancer, the Merkel Cell Carcinoma (MCC). Although MCPyV is constantly shed from healthy skin, MCC incidence increases among aging and immunocompromised individuals. To date, the events connecting initial MCPyV infection and subsequent transformation still remain elusive. MCPyV differs from other known polyomaviruses concerning its cell tropism, entry receptor requirements, and infection kinetics. In this study, we examined the cellular requirements for endocytic entry as well as the subcellular localization of incoming virus particles. A thorough understanding of the determinants of the infectious entry pathway and the specific biological niche will benefit prevention of virus-derived cancers such as MCC.

## Introduction

Polyomaviruses (PyV) are small, non-enveloped dsDNA viruses with a diameter of 45-50 nm. The icosahedral (T=7) capids consist of 72 homopentameric capsomers of the major capid protein VP1 with minor capsid proteins VP2/VP3 located within a cavity underneath the VP1 pentamers. The PyV capsid harbors a chromatinized, circular dsDNA genome of about 5 kb (1, 2). Well-studied PyV such as simian virus 40 (SV40), and murine PyV (mPyV) possess a broad cell tropism and can transform cells *in vitro* and in animals (3, 4). Of the human polyomaviruses, JC and BK virus are the best studied (5, 6). JC and BK viruses were initially identified in brain and urine samples, respectively (7, 8). Initial infection with these viruses occurs early in life and typically leads to persistent infections that are typically benign (9-11). Upon immunosuppression, however, persistent JC and BK virus infections may lead to severe diseases, such as progressive multifocal leukoencephalopathy or PyV-associated nephropathy, with potentially fatal outcomes (3, 4).

In 2008, Feng and colleagues identified Merkel cell polyomavirus (MCPyV) in a rare form of skin cancer known as Merkel cell carcinoma (MCC) (12). MCC is an aggressive cancer with increasing incidence (13, 14), which is most likely to develop in immunocompromised and elderly populations upon prolonged UV exposure (15, 16). About 80% of MCC are positive for MCPyV DNA integrated into the host genome (12). As for most PyV, MCPyV infections are widespread and predominantly asymptomatic. In fact, MCPyV is continuously shed from healthy skin with a prevalence of 60-80% (17-19). However, MCPyV DNA has also been isolated from respiratory, urine and blood samples (20), and the range of tissues in which persistent infection can be established is thus still unclear. The presence of integrated MCPyV DNA in MCC cells is thought to cause cancer through the continuous expression of the transforming large T (LT) and small T (sT) antigens. Integration of the viral DNA into the host cell genome is coupled to truncation of the LT C-terminal domain, which is important for viral genome replication and can induce p53 activity, triggering cell cycle arrest (21, 22). The viability of MCC cells depends on expression of LT and/or sT, as pan-T knock-down in MCC-derived cells lead to cell death (23, 24). Importantly, the cellular origin of MCC is still under debate. A recent report suggests dermal fibroblasts as target cells for productive infection, whereas Merkel cells are not permissive for virus entry or productive infection (25, 26). Thus, it remains unclear exactly which events give rise to MCC.

Since cell culture systems to produce sufficient quantities of infectious MCPyV are not readily available, MCPyV vectors, so called pseudoviruses (PsV), are important tools to study entry. As MCPyV does not contain detectable levels of VP3 (27), PsV consist of VP1/VP2-only capsids that harbor a reporter plasmid (e.g. coding for EGFP or luciferase). Expression of the reporter allows easy readout for successful entry, i.e. delivery of the viral DNA to the site of transcription and replication (28). In an effort to identify MCPyV-permissive cell lines and to better understand the tissue tropism of MCPyV, the tumor cell library NCI-60 was screened for transducibility and ability to support virus replication with MCPyV PsV and native virions, respectively (29). Of those, A549 cells, a non-small cell lung cancer cell line, showed robust transducibility with MCPyV PsV (28).

Since MCPyV is an emerging virus, little is known about the basic biology of the virus, in particular how initial infection occurs. Initial studies on the mechanism of MCPyV infection addressed cell surface interactions and cellular tropism of MCPyV (25, 28, 30). MCPyV relies on binding sulfated glycosaminoglycans (GAGs) for initial attachment similar to papillomaviruses (28). However, it also requires interaction with carbohydrates containing a linear siali acid motif, i.e. Neu5Ac□2-3Gal, like other PyV (28, 30, 31).

Different viruses utilize distinct preexisting cellular endocytosis pathways for infectious entry (32). These endocytic pathways are characterized by a specific set of cellular factors and regulators facilitating endocytic vesicle formation. This set of cellular factors is used to distinguish different pathways. Well-studied endocytic pathways include clathrin-mediated endocytosis (CME), caveolar/lipid raft-mediated endocytosis, and macropinocytosis (33), which are essential for receptor-mediated endocytosis, turnover of plasma membrane receptors and fluid-phase uptake. To facilitate safe delivery and release of the viral genome to and at the site of replication, viruses are routed through specific intracellular target organelles, such as endosomes, the Golgi apparatus, or the endoplasmic reticulum (ER).

Host cell entry of most PyV occurs by caveolar/lipid raft-mediated endocytosis (34-37), whereas JC virus, for example, uses CME (38). After endocytosis, virions are routed through the endosomal pathway to be delivered to the ER (39). There, a chaperone- and disulfide isomerase-mediated uncoating step occurs, whereupon the modified particles are translocated into the cytosol by the ER-associated degradation machinery (40-43). In the current study, we aimed to identify the cellular requirements for virus entry into host cells to understand similarities and differences to other PyV. For this, we used small compound inhibitors of decisive cellular factors/processes during MCPyV infection. As functional controls, we employed well-studied viruses, including SV40, the papillomavirus HPV16, or the alphavirus Semliki Forest Virus (SFV) (36, 44, 45). In addition, we characterized the entry route of MCPyV by ultrathin section transmission electron microscopy (TEM). Our results support a model in which MCPyV enters A549 cells by caveolar/lipid-raft dependent endocytosis, and where viruses are routed through the endosomal pathway to the ER. To our surprise, we found that MCPyV acquired a lipid membrane within endosomes. This membrane was absent once the virus located in the ER.

## Results

### MCPyV PsV cell binding and infection depend on sulfated glycosaminoglycans

To study the infectious entry requirements of MCPyV, we tested the susceptibility of the keratinocyte-derived cell lines HeLa and HaCaT in comparison to A549 cells. Using PsV we found that neither HeLa nor HaCaT cells showed GFP expression 72 h post infection and are thus poorly permissive for MCPyV entry (Fig. 1 A). We, therefore, decided to use A459 cells for our study, which allow for infectious internalization as previously described (29).

**Figure 1:**
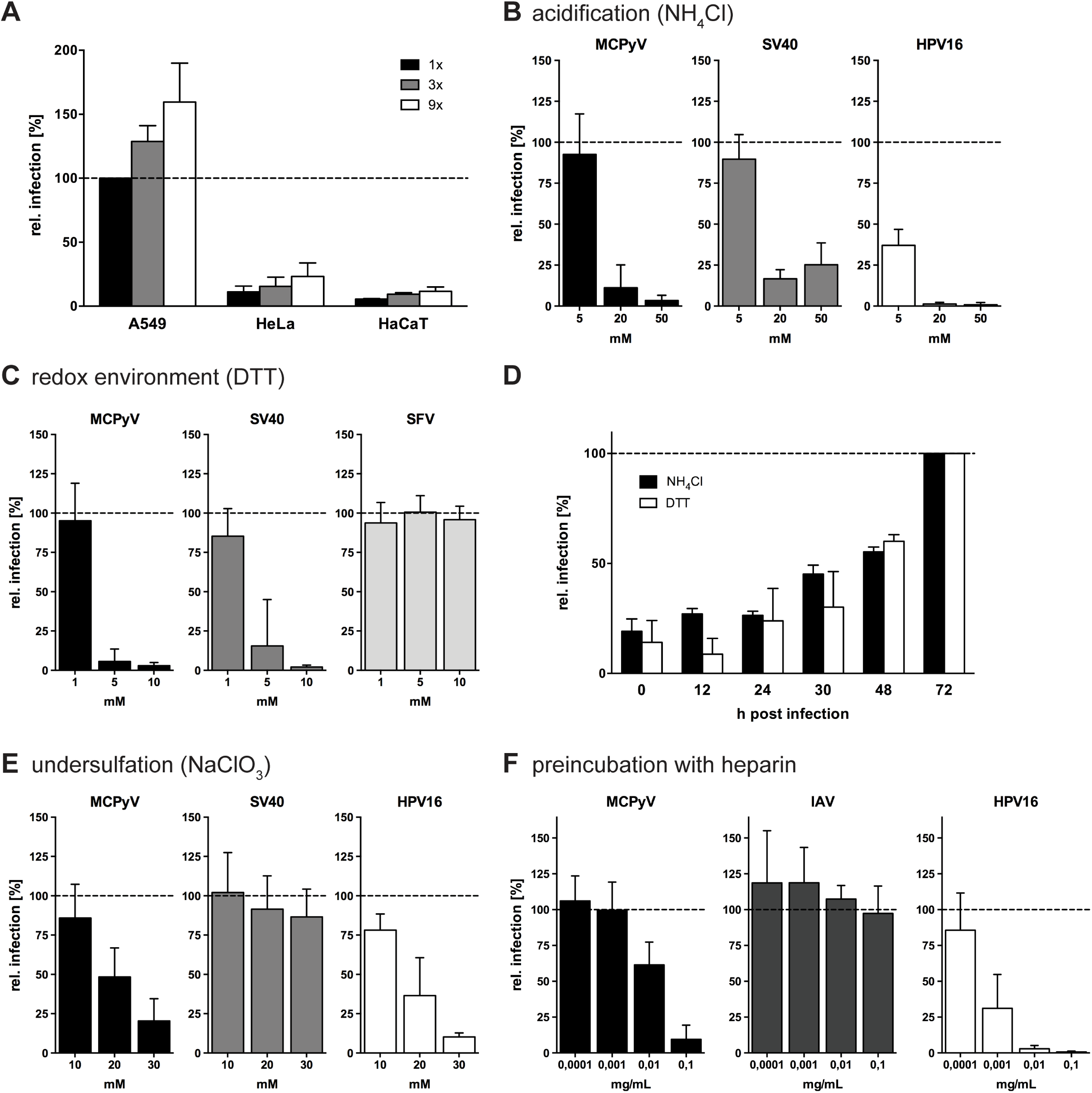
MCPyV infection is slow and asynchronous, and relies on interaction with sulfated glycans. (A) A549, HeLa or HaCaT cells were infected with increasing amounts of MCPyV PsV. Depicted are relative infection values (%) related to the MCPyV amount yielding 20% absolute infection in A549 cells ± SD (1x = 80 ng VP1). (B) A549, CV-1 or HeLa cells were infected with MCPyV, SV40 or HPV16, respectively, in the presence of indicated concentrations of NH4Cl. (C) A549, CV-1 or BHK cells were infected with MCPyV, SV40 or SFV, respectively, in the presence of indicated concentrations of DTT. (D) A549 cells were infected with MCPyV for total 72 h, while 10 mM NH_4_Cl or 5 mM DTT was added at indicated h p.i.. (E) A549, CV-1 or HeLa cells were infected with MCPyV, SV40 or HPV16, respectively, upon pretreatment with indicated concentrations of NaClO_3_ for 16 h prior to infection. (F) A549 were infected with MCPyV or IAV, and HeLa cells were infected with HPV16 virions, which were treated with indicated concentrations of heparin for 1 h prior to infection. (B-F) Depicted are averages of relative infection values to untreated controls ± SD from at least 3 independent experiments.

As initial strategy to elucidate the cellular route(s) of entry, perturbation of cellular processes followed by infection is key to identify factors/organelles facilitating MCPyV entry. The time course of MCPyV infectious entry into A549 cells was rather slow and plateaued after 72 h post infection (p.i.) (data not shown). Since indirect effects occur often after prolonged cellular perturbation, it is important to minimize such effects (46). For this, we compared siRNA-mediated knockdown, expression of dominant negative mutants, and small compound inhibition (data not shown). The efficacy and reliability of siRNA-mediated knockdown and expression of dominant mutants turned out to be questionable, as judged by infection of control viruses, so that we turned exclusively to small compound inhibition.

We first tested whether inhibition of acidification of endosomal compartments with the weak base ammonium chloride (NH_4_Cl) perturbed MCPyV infection (47). In the presence of NH_4_Cl, MCPyV infection was blocked in a dose-dependent fashion (Fig. 1 B) similar to infection with SV40 and HPV16, which served as positive controls (45, 48). This indicates the requirement for low pH in endosomal compartments during infectious internalization of MCPyV.

SV40 is directed to the ER for uncoating and subsequent translocation to the cytosol using ER resident disulfide oxidoreductases and the ER-associated degradation machinery (40). Dithiothreitol (DTT) is a powerful reducing agent and thus perturbs the cellular redox environment and disulfide oxidoreductase functions (49). Accordingly, DTT treatment arrests SV40 in the ER (40). We tested whether MCPyV was also sensitive to DTT treatment. MCPyV and SV40 infectivity was clearly reduced in the presence of DTT, whereas entry of Semiliki Forrest virus (SFV), which does not require a redox-driven ER uncoating step (44), was unaffected (Fig. 1 C).

Next, the ability of NH4Cl or DTT to block MCPyV infection in relation to the time course of entry was tested. For this, the drugs were added different times p.i.. NH_4_Cl and DTT exhibited a similar propensity to block MCPyV infection, where infectivity was inhibited upon early addition and increased with additions at later time point (Fig. 1 D). The increase at later time points indicates that the virus had passed the step blocked by the drugs. At 30 h p.i., about 50% of MCPyV had passed the acid-activated step, whereas the redox-dependent step appeared to occur slightly later (Fig. 1 D). The time courses indicate in addition a rather slow and asynchronous entry process akin to HPV16 (Fig. 1 D, (45)).

Prolonged treatment of cells with drugs that block endocytosis and trafficking can have indirect or cytotoxic effects. To address this problem, we employed an inhibitor swap approach in which cells were transiently treated with a drug of interest followed by replacement by another that blocks entry at a later stage, such as acid activation in endosomes or redox-mediated uncoating in the ER (45, 46). Moreover, several different viruses served as positive and negative controls to assess the efficacy of treatment as well as the potential for pleiotropic drug effects.

Initially, as proof of feasibility, the requirement for sulfated GAGs was tested. MCPyV attachment to cells generally depends on such GAGs (28, 30, 31). Therefore, treatment with sodium chlorate (NaClO_3_) was used to induce production of undersulfated GAGs (50). Treatment of A549 cells with 10, 20, or 30 mM NaClO_3_ for 16 h prior to inoculation with MCPyV produced a dose-dependent reduction in infectivity (Fig. 1 E). HPV16 infectivity was similarly reduced, as expected (51). In contrast, infection with SV40 was unaffected, suggesting different requirements for the two PyV (Fig. 1 E). In addition, we tested the inhibitory effect of heparin, a highly sulfated GAG, on MCPyV infection. In line with previous results (28), preincubation of MCPyV pseudovirions with heparin efficiently blocked infection, only slightly less efficiently than the positive control HPV16 (Fig. 1 F). Influenza A virus (IAV) was used as a negative control, as IAV infection depends on interaction with sialic acids (52) and is independent of GAG binding. Heparin did not affect IAV infectivity (Fig. 1 F). Taken together, the results confirm that MCPyV infection requires interactions with sulfated cell-surface GAGs.

### Inhibitory profile of MCPyV endocytosis

After confirming previous reports, we set out to characterize the endocytic pathway employed by MCPyV. First, CME was blocked with the small molecule inhibitor pitstop2, which interacts with the clathrin N-terminal domain and thereby interferes with assembly of the clathrin coat (53). Infection with MCPyV was unaffected by treatment with pitstop2 (Fig. 2 A), similar to SV40, which enters cells by caveolar/lipid-raft endocytosis (34, 36, 48). As expected, infection with SFV, which enters cells by CME (44, 54), was efficiently reduced. Thus, MCPyV entry occurs independently of CME.

**Figure 2:**
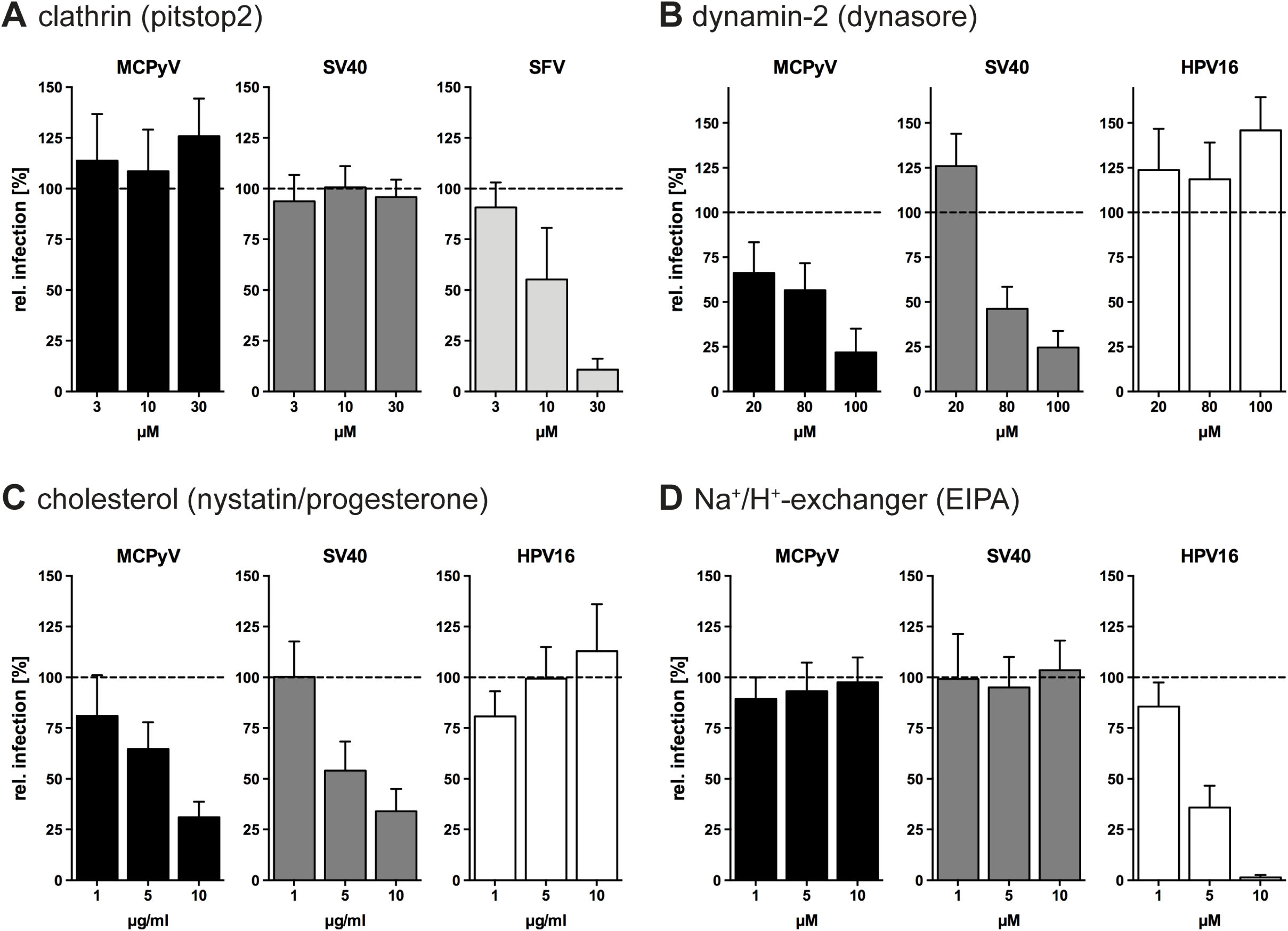
MCPyV infection is dynamin- and cholesterol-dependent, but independent from clathrin and Na^+^/H^+^-exchangers. (A) A549, CV-1 or BHK cells were infected with MCPyV, SV40 or SFV, respectively, in the presence of indicated concentrations of pitstop2. (B-D) A549, CV-1 or HeLa cells were infected with MCPyV, SV40 or HPV16, respectively, in the presence of indicated concentrations of dynasore (B), nystatin/progesterone (C), and EIPA (D). Depicted are percentages of infection values relative to solvent treated controls ± SD of at least 3 independent experiments.

Another important regulator of several endocytic pathways is dynamin-2 (Dyn2). The large GTPase regulates scission of endocytic pits in several endocytic pathways, i.e. CME, caveolae/lipid raft-mediated endocytosis, IL-2 endocytosis, and phagocytosis (32). To study a potential Dyn2 involvement in MCPyV uptake, dynasore (an inhibitor of the dynamin GTPase activity) was employed (55). SV40 served as a positive control as it is known to depend on Dyn2 activity (56). Infection of MCPyV was blocked dose-dependently by dynasore to a residual level of 22±13%, similar to SV40 (Fig. 2 B). As a negative control, HPV16 was used. HPV16 enters host cells by a clathrin-, caveolin-, cholesterol- and dynamin-independent, but actin-dependent endocytic pathway (45). As expected, HPV16 infection remained unperturbed or even increased after dynasore treatment (Fig. 2 B). This indicates that MCPyV is endocytosed by a Dyn2-dependent pathway similar to SV40 but distinct from HPV16 endocytosis.

To further ascertain that MCPyV enters cells by a similar endocytic mechanism as SV40, we perturbed cholesterol-dependent caveolar/lipid-raft endocytosis. To this end, we treated cells with nystatin and progesterone, which sequester cholesterol and prevent cholesterol synthesis, respectively (34, 56-59). As expected, infection with MCPyV and SV40 were inhibited upon perturbation of cholesterol to residual levels of 31±8% and 34±11%, respectively (Fig. 2 C), whereas HPV16 infection remained unaffected as described previously (45). This suggests that MCPyV infection, like SV40 infection, requires cholesterol-rich membrane domains (34).

Next, a potential role for macropinocytosis was assessed. The Na^+^/H^+^-exchanger regulates macropinocytosis by controlling submembraneous pH, which in turn regulates the formation of membrane protrusions (60). Ethylisopropylamiloride (EIPA)-mediated inhibition of the Na^+^/H^+^-exchanger (61) is a classical treatment to interfere with macropinocytosis. This treatment neither affected MCPyV nor SV40 infection (Fig. 2 D), whereas HPV16 infection was efficiently blocked as expected (45). Thus, MCPyV endocytosis occurs independently of the Na^+^/H^+^-exchanger and, consequently, also of macropinocytosis or related pathways.

In summary, MCPyV entry required cholesterol-rich membranes and dynamin, but did not require clathrin or macropinocytic pathways. These observations are most consistent with an entry pathway that employs caveolar/lipid-raft mediated endocytosis.

### Regulation of MCPyV endocytosis depends on actin dynamics, Rho-like GTPases and signaling via Tyr kinases and phosphatases PP1 and PP2A/B

Next, we addressed functional regulators involved in endocytic processes with respect to their role in MCPyV entry.

Actin polymerization is required for most endocytic pathways employed by viruses, where formation of protrusions, intracellular transport or vesicle scission is facilitated (32, 45). Cytochalasin D and jasplakinolide inhibit actin dynamics by either blocking polymerization or depolymerization, respectively (62, 63). Both cytochalasin D and jasplakinolide efficiently blocked MCPyV and SV40 infection (Fig. 3 A, B), where the inhibitory effect of jasplakinolide was less pronounced for SV40. In contrast, CME-mediated uptake of SFV was unaffected by stabilization of actin filaments (Fig. 3 B) and increased upon actin depolymerization (Fig. 3 A).

**Figure 3:**
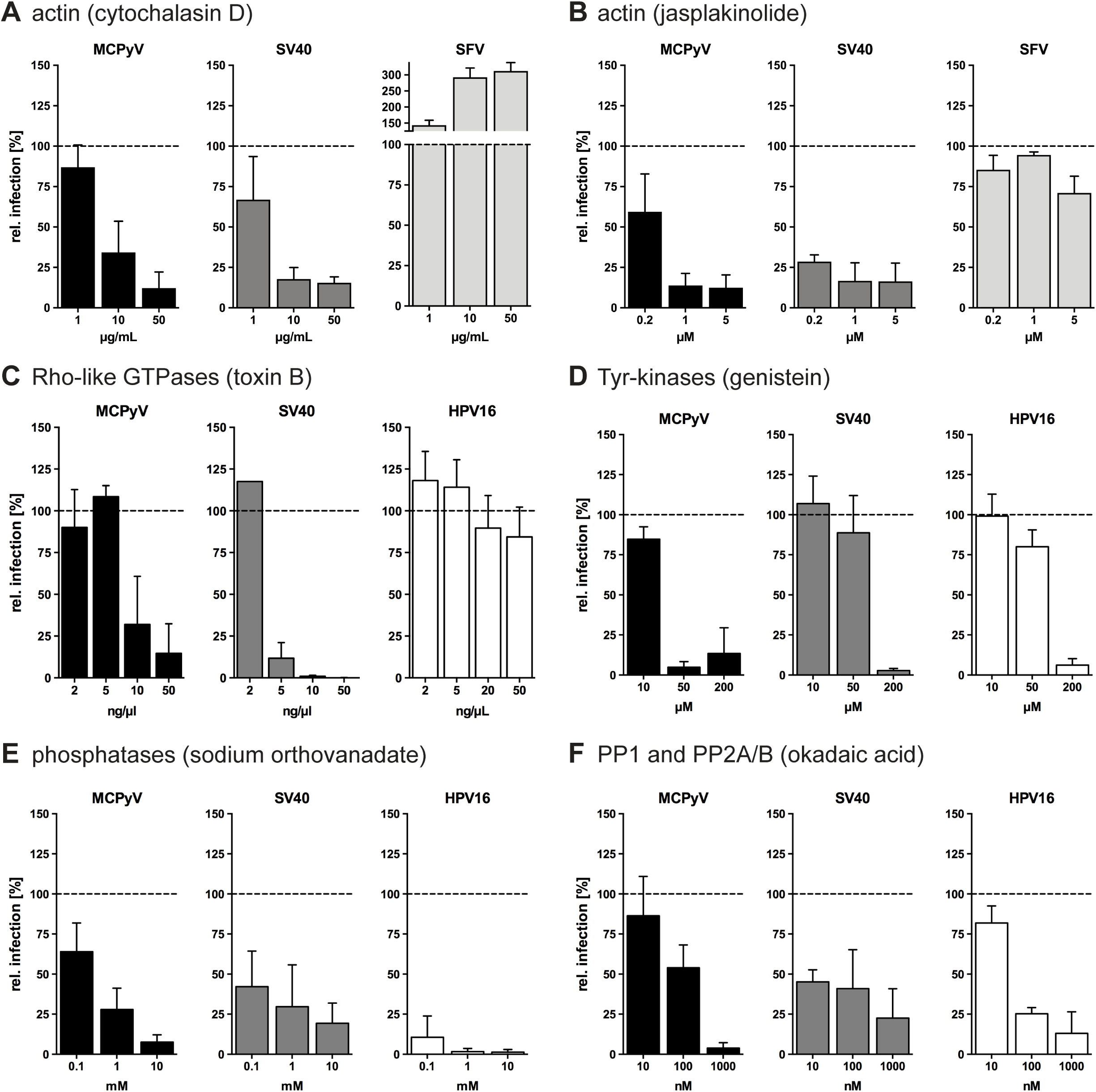
MCPyV infection is dependent on actin, RhoGTPases, tyr-kinases, and cellular phosphatases. (A-B) A549, CV-1 or BHK cells were infected with MCPyV, SV40 or SFV, respectively, in the presence of indicated concentrations of cytochalasin D (A) or jasplakinolide (B). (C-F) A549, CV-1 or HeLa cells were infected with MCPyV, SV40 or HPV16, respectively, in the presence of indicated concentrations of toxin B (C), genistein (D), sodium orthovanadate (E), or okadaic acid (F). Depicted are percentages of infection values relative to solvent treated controls ± SD of at least 3 independent experiments.

Actin polymerization is most often regulated by the activity of pathway-specific Rho-like GTPases (32, 64, 65). Toxin B from *C. difficile* is a broad inhibitor of all Rho-like GTPases (66) and was hence used to assess the relevance of Rho GTPases for infection by MCPyV and SV40. As a negative control, we used HPV16 infection, which was not perturbed by treatment with toxin B, as expected (Fig. 3 C, (45)). Toxin B reduced MCPyV and SV40 infection to 15±18% and to 0.1±0.1% residual infection, respectively (Fig. 3 C). From this we conclude that Rho-like GTPases likely mediate actin-dependent steps during MCPyV endocytosis (Fig. 3 C). Analogous to SV40, the key step may be actin-dependent closure and fission of endocytic vesicles from the plasma membrane (56).

Ligand-induced activation of tyrosine kinases (tyr-kinases) regulates the activation of several endocytic pathways such as macropinocytosis and caveolar/lipid raft-mediated endocytosis, whereas CME and other pathways are tyr-kinase independent (32, 65). Genistein, a broad and efficient inhibitor of tyr-kinases (67), was used during infection with MCPyV, SV40, and HPV16. As expected for HPV16 and SV40, infection was blocked upon treatment with 200 µM genistein (Fig. 3 D). Interestingly, MCPyV infection was already sensitive to 50 µM genistein treatment (Fig. 3 D) suggesting a strong requirement for tyr-kinases, potentially at multiple levels.

Next, cellular phosphatases were inhibited with the broadly active sodium orthovanadate, a competitive inhibitor of all phosphatases (68, 69). Entry of MCPyV, SV40 and HPV16 were all strongly inhibited in the presence of orthovanadate (Fig. 3 E). Okadaic acid, a more specific inhibitor of phosphatases families PP1, PP2A and PP2B (70), also inhibited MCPyV, SV40 and HPV16 (Fig. 3 F).

In summary, MCPyV entry is likely facilitated by dynamic actin rearrangement that is regulated by Rho-like GTPases, as well as by the activity of tyr-kinases and PP1 and/or PP2A/B phosphatases.

### Intracellular trafficking of MCPyV requires endosomal acidification, functional microtubular dynamics, and an intact redox environment

Treatment with NH_4_Cl reduced MCPyV infection indicating an acid-activation step (Fig. 1 B). To test whether this acid-activated step occurred in endosomes, bafilomycin A1, an inhibitor of the endosomal proton pump V-ATPase, was used (71). Bafilomycin A1, like NH_4_Cl, inhibited the infectivity of MCPyV, SV40, and HPV16 in a dose-dependent fashion (Fig. 4 A). Thus, MCPyV depends on low endosomal pH for either acid activation or trafficking within maturing endosomes.

**Figure 4:**
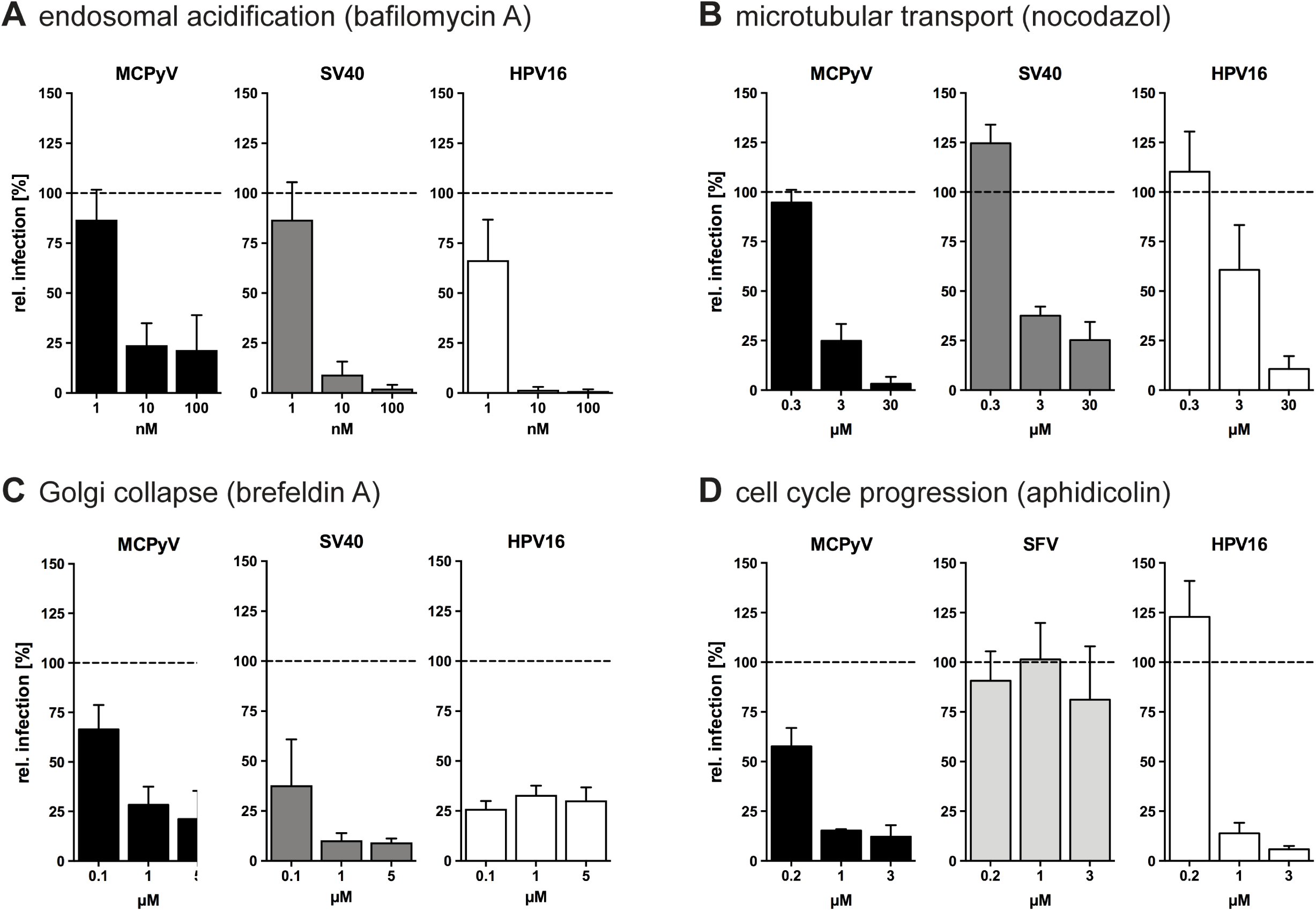
MCPyV infection requires endosomal acidification, functional microtubular dynamics, and an intact redox environment. (A-C) A549, CV-1 or HeLa cells were infected with MCPyV, SV40 or HPV16, respectively, in the presence of indicated concentrations of bafilomycin A (A), nocodazol (B), and brefeldin A (C). A549, BHK, or HeLa cells were infected with MCPyV, SFV or HPV16 respectively, in the presence of indicated concentrations of aphidicolin (D). Depicted are percentages of infection values relative to solvent treated controls ± SD of at least 3 independent experiments.

Most intracellular transport occurs along microtubules (72). To assess microtubule involvement in MCPyV infection, depolymerization of microtubules by treatment with the polymerization blocker nocodazole was used (73). As reported previously, SV40 and HPV16 entry require the integrity of microtubules (Fig. 4 B). Similarly, MCPyV infection strongly depended on intact microtubular dynamics (Fig. 4 B). This suggests that MCPyV requires microtubules for transport of vesicular compartments or of the virus itself during host cell entry.

To study the involvement of intracellular transport processes from the ER to the Golgi apparatus in MCPyV entry, we used brefeldin A, which eventually leads to Golgi collapse into the ER (74). Upon perturbation with brefeldin A, infection with MCPyV, SV40, and HPV16 were blocked (Fig. 4 C). This indicates that infection with these viruses requires the functional integrity of the secretory ER/Golgi compartments.

Another hint for an involvement of the ER during infectious internalization can be drawn from the perturbation of the cellular redox environment by DTT. At the concentrations used, DTT interferes mostly with the formation of disulfide bonds during folding in the ER but not with existing disulfide bonds in folded proteins (49). Since MCPyV infection was blocked by DTT (Fig. 1 C), it is reasonable to assume that MCPyV like SV40 requires an ER step for host cell entry, possibly for uncoating and translocation into the cytosol.

After escape from the ER, SV40 is thought to be imported into the nucleus via the nuclear pore complexes (75). In contrast, HPV16 requires cell cycle progression, i.e. nuclear envelope breakdown, for nuclear entry (76, 77). To address whether MCPyV infection depends on mitotic activity for nuclear entry, we blocked cells in S-phase using aphidicolin, an inhibitor of DNA polymerase alpha and delta (78). MCPyV entry was efficiently blocked by aphidicolin to a similar extent as HPV16 (Fig. 4 D). The RNA virus SFV replicates in the cytoplasm and therefore served as a negative control (79). SFV infection was not affected by aphidicolin treatment (Fig. 4 D), as expected. These results indicate that MCPyV infection may depend on the mitotic activity of the host cells.

### MCPyV is internalized into small, non-coated vesicles

Our inhibition studies indicate that MCPyV and SV40 use similar pathways to infect cells. To confirm this notion, we examined virion trafficking using TEM. Localization of virus particles was assessed 2, 8, 16, 24 and 48 h p.i.. Interestingly, the virus localized to several specific cellular compartments at each time-point, in line with asynchronous internalization and trafficking of the MCPyV particles. Virus particles were readily detectable bound to the cell surface (Fig. 5 A). In addition, they were found within small, inward budding membrane invagination without any visible protein coat (Fig. 5 A, B). Interestingly, virions resided in two distinct but equally frequent populations of invaginations. In one class of invagination, the plasma membrane appeared to make close contact with the virion surface, while the second class of invagination exhibited 10 nm ± 8 nm distances between the membrane and the virion surface. The first class of invagination is reminiscent of prior observations for SV40, which is known to make close contact with the sialylated headgroups of plasma membrane glycolipids (80, 81). The latter class of invagination suggests possible interactions between the virion and a glycoprotein receptor with a large ectodomain, such as an HSPG. In addition, MCPyV virions were found within intracellular vesicles without an apparent protein coat (Fig. 5 C). These data, and the absence of virus in large, coated or tubular invaginations, is consistent with MCPyV uptake by caveolar/lipid raft-mediated endocytosis rather than uptake by macropinocytosis, CME or glycosphingolipid-enriched carriers (58, 81-83).

**Figure 5:**
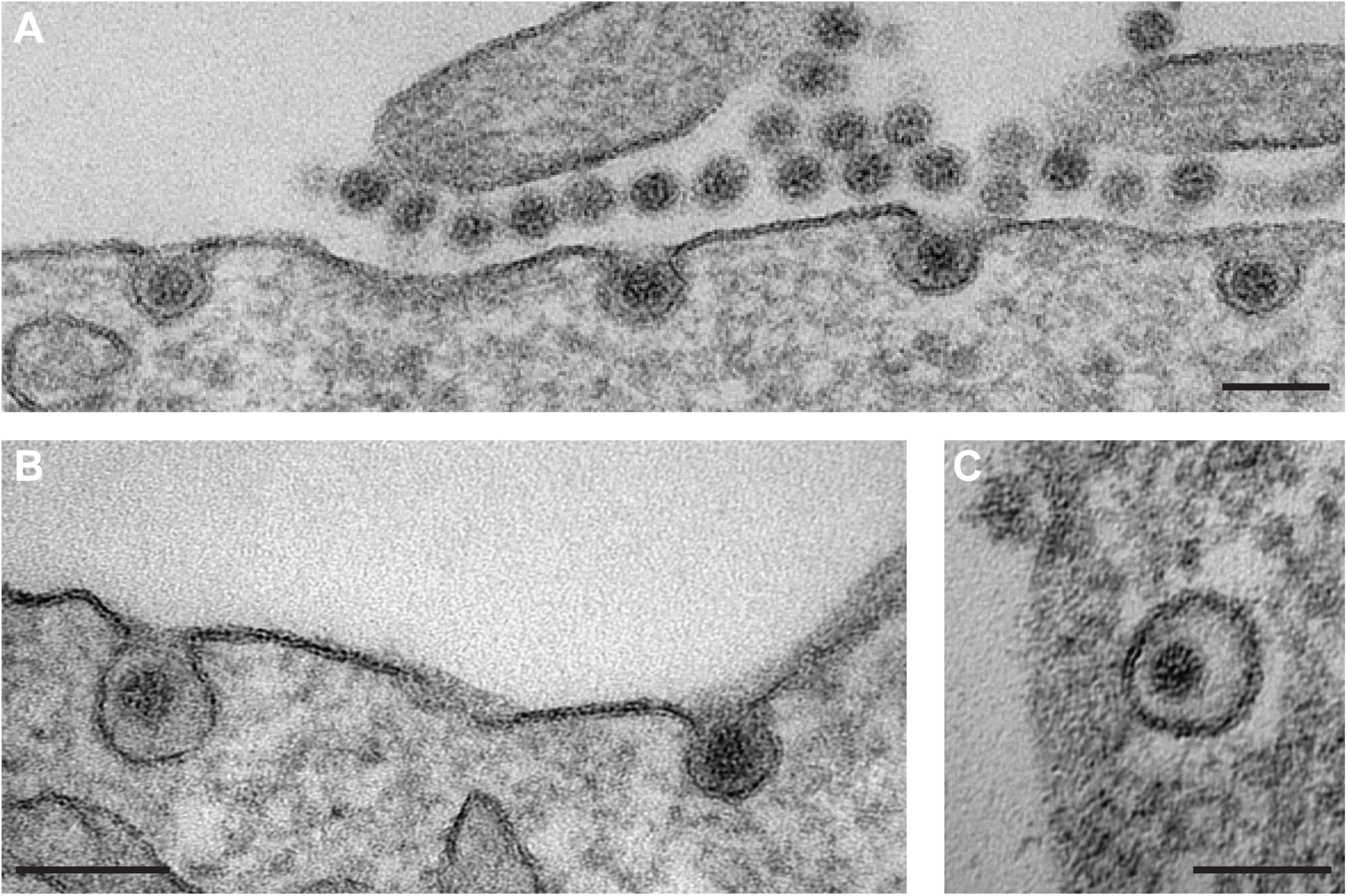
MCPyV is taken up into tight-fitting inward budding pits. (A-C) A549 cells were infected with MCPyV particles for 2-24 h before fixation with glutaraldehyde. Cells were processed for TEM according to standard procedures. Images of early entry events were taken. Scale bar 100 nm.

### MCPyV traffics via the endolysosomal route to the ER

After initial uptake into endocytic vesicles, MCPyV particles were found in early endosomal compartments and later accumulated in late endosomal/lysosomal structures (Fig. 6 A, B). Other polyomaviruses, such as SV40, JCPyV and BKPyV, are then trafficked to the ER via Golgi- and non-Golgi routes. In the ER, oxidoreductases and chaperones mediate partial disassembly of the virions (uncoating) and transfer of partially disassembled virions into the cytosol (40, 81, 84-86). MCPyV was undetectable in the Golgi apparatus (Fig. 6 D), the cytosol, or the nucleus. Similar to other PyV, MCPyV was observed in the lumen of the ER, but only to a minor extent (Fig. 6 C).

**Figure 6:**
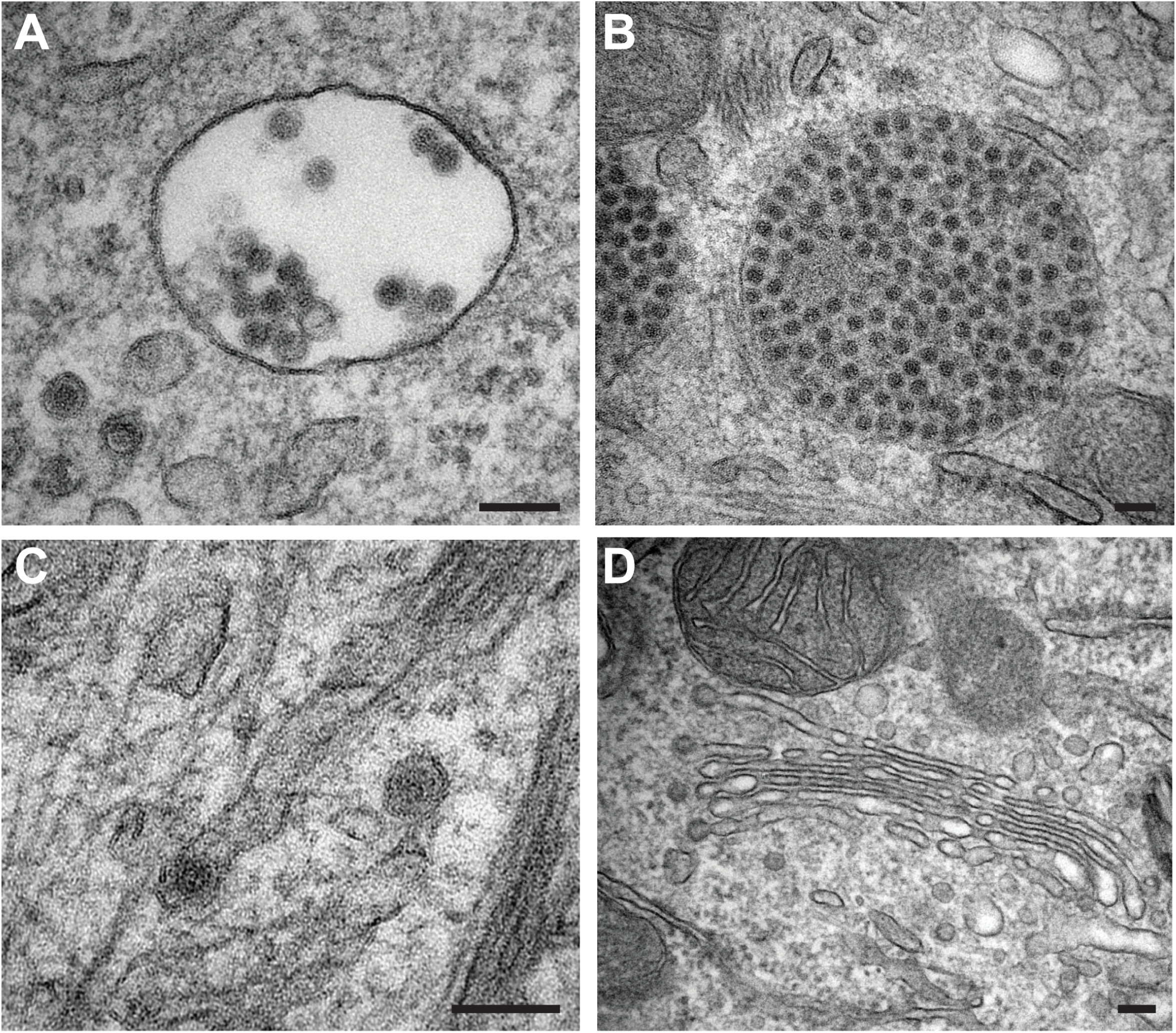
MCPyV travels through the endolysosomal system to the ER omitting the Golgi. (A-D) A549 cells were infected with MCPyV particles for 2-24 h before fixation with glutaraldehyde. Cells were processed for TEM according to standard procedures. Virus particles were found in endosomal compartments (A, B) and in the ER (C). MCPyV was absent from the Golgi (D). Scale bar 100 nm.

Taken together, the EM results thus support the conclusion that MCPyV and SV40 use similar infectious entry pathways.

### MCPyV acquires a lipid envelope in endosomal compartments

In addition to the observations described above, we surprisingly found MCPyV particles with what appeared to be a double layer lipid membrane in endosomal compartments (Fig. 7 A, B). The membrane tightly enveloped the particles. The enveloped virions were easily detectable in maturing and late endosomes as well as endolysosomes similar to non-enveloped particles. In fact, both enveloped and non-enveloped particles were detectable side by side in the same endosomes (Fig. 7 C, enveloped black arrows vs. non-enveloped white arrow). This suggested that the process of acquiring a lipid envelope occurs during viral passage of the endosomal pathway rather than in distinct, specialized organelles. It is unclear whether the enveloped particles are part of the productive infectious entry pathway or are instead a dead end for the virus. Possibly, this phenotype reflects a new mode of antiviral defense or a mechanism to evade endosomal degradation.

**Figure 7:**
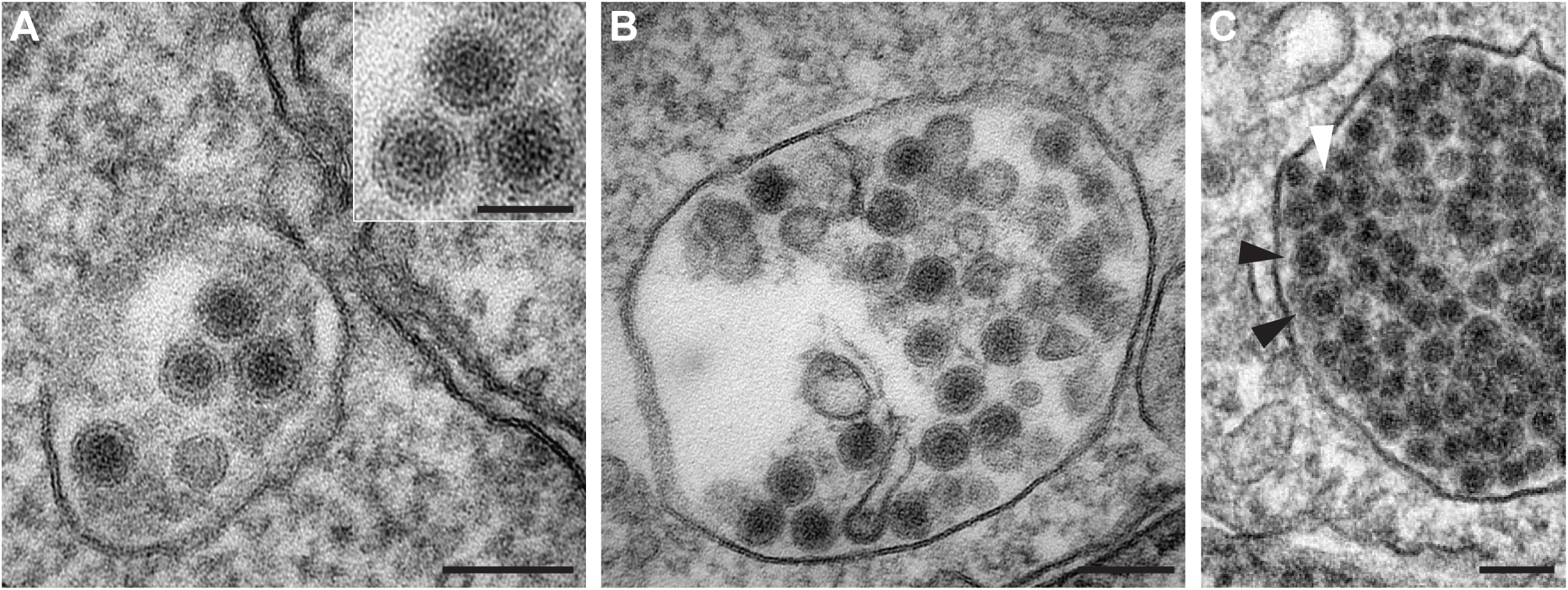
MCPyV virions acquire a membrane envelope during cell entry. (A-C) A549 cells were infected with wild-type MCPyV particles before fixation with glutaraldehyde. Cells were processed for TEM according to standard procedures. Images of virus particles in intracellular compartments were taken after 8 h and 16 h p.i.. Note the envelope around the virus particles in the representative endosomal compartments. C) Note that enveloped particles (black arrows) are found side-by-side with non-enveloped virions (white arrow) in the same organelle. Scale bars 100 nm; Scale bar inset 50 nm.

### The role of sialic acid for MCPyV entry

Previous studies showed that mutations in the sialic acid binding pocket of MCPyV VP1 rendered the particle non-infectious but did not perturb binding to cells (30). Hence, sialic acid interaction is essential for infectious entry, whereas initial attachment occurs mainly by GAG engagement. Since we observed MCPyV particles in endocytic pits in two distinct populations that may reflect binding to gangliosides and HSPGs, two sialic acid binding site mutants (W76A and Y81V) were followed using EM. The mutant PsV assembled without visible defects (Fig. 8 A, B i, C i), but were unable to mediate infection in A549 cells (Fig. 8 D), as expected (30). To verify the desired glycan binding abilities of the sialic acid mutants, we conducted saturation transfer difference (STD-) NMR experiments to probe for binding of sialylated oligosaccharidsses and GAG oligosaccharides to MCPyV wild-type and mutant virus-like particles (VLPs). STD-NMR makes use of energy transfer from proteins to their ligands upon binding and thus allows determination of binding specificities by comparing the changes in the resonance frequencies of the ligand molecules in association with wild-type or mutant viruses (30, 87). Here, 3’sialyllactose (3’SL) and a chemically well-defined heparan sulfate (HS)- pentasaccharide, Arixtra (Ax), were chosen as a minimal sialylated MCPyV ligand and a short GAG oligosaccharide, respectively (Fig. 9 A, B). STD-NMR showed that the GAG pentasaccharide bound to wild-type VLPs as well as to the W76A mutant (Fig. 9 B ii vs. iii). In contrast, 3’SL bound both wild-type and Y81V mutant VLPs but no saturation transfer was observed from the W76A mutant to 3’SL (Fig. 9 A ii vs. iii vs. iv). Consistent with previous crystallographic and NMR-based results (30), only resonances from the 3’SL sialic acid and galactose rings were observed. For the HS pentasaccharide (Fig. 9 B), on the other hand, magnetization transfer was observed almost equally for all five monosaccharide units, with the non-sulfated GlcA ring being the only unit, whose resonances were somewhat underrepresented in the STD-NMR spectra in comparison to the ^1^H reference spectrum of the free pentasaccharide. This suggests that the GlcA ring is the least important determinant in the interaction. In summary, these results confirm that the W76A mutant is defective in sialic acid binding, whereas Y81V remained partially able to interact with the 3’SL ligand under these conditions. Furthermore, the binding site for GAGs apparently remained intact upon mutation suggesting that the GAG binding site is spatially distinct from the sialic acid interaction site.

**Figure 8:**
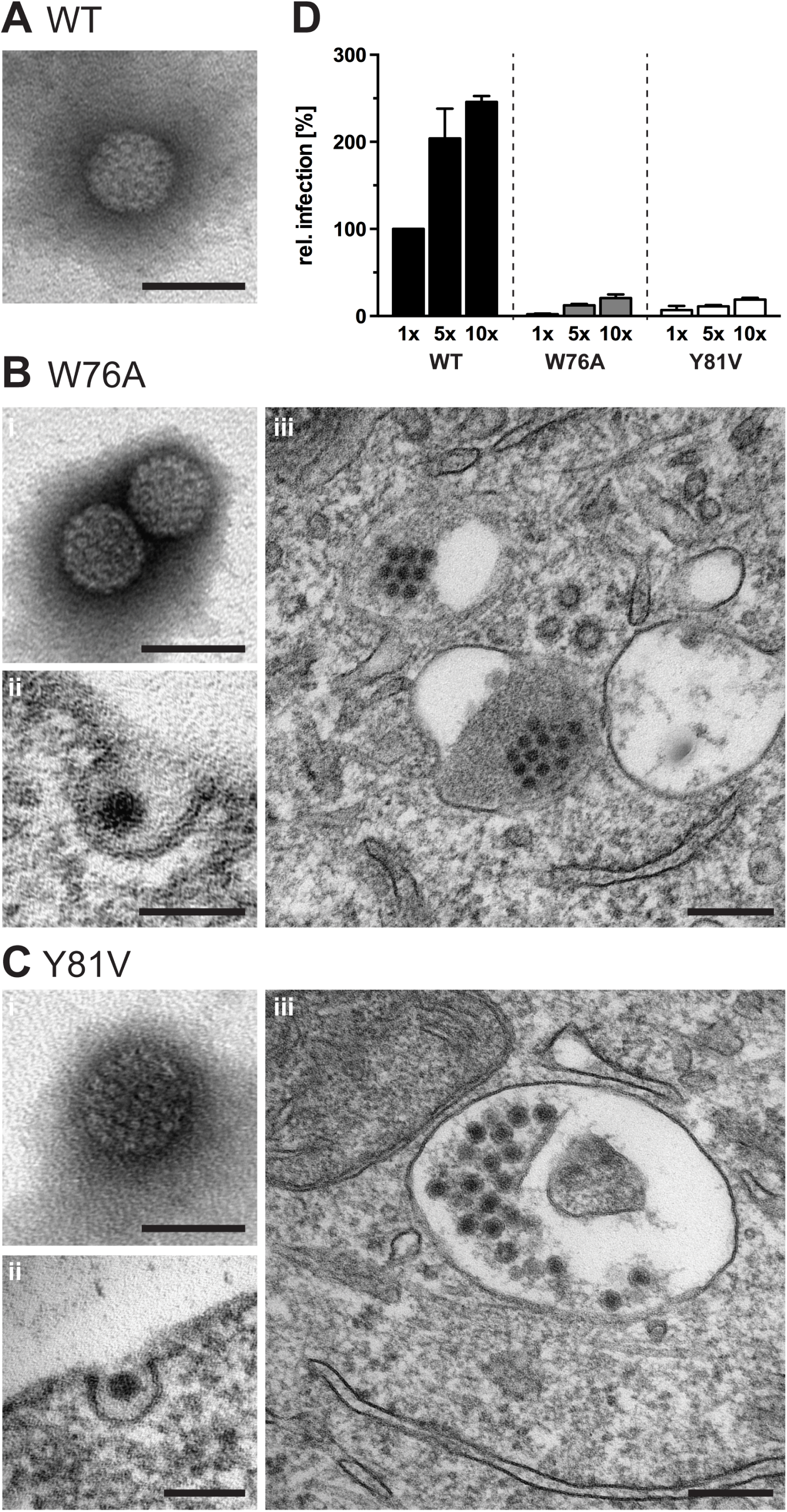
MCPyV sialic acid binding mutants are taken up into A549 cells but fail to mediate infection. (A-C i) MCPyV wild-type and mutant particles were analyzed by electron microscopy after negative staining. Depicted are representative images of virions. Scale bars: 50 nm. (B-C ii and iii) A549 cells were infected with W76A and Y81V MCPyV particles for 2-24 h before fixation with glutaraldehyde. Cells were processed for TEM according to standard procedures. Images of virus particles in endocytic pits and in intracellular compartments were taken after 8 h (ii) and 16 h p.i. (iii), respectively. Scale bar 100 nm for ii and 200 nm for iii. (D) 60 ng (1x), 300 ng (5x) and 600 ng (10x) of MCPyV wild-type and W76A and Y81V mutant particles were used in an infection assay with A549 cells for 72 h.

**Figure 9:**
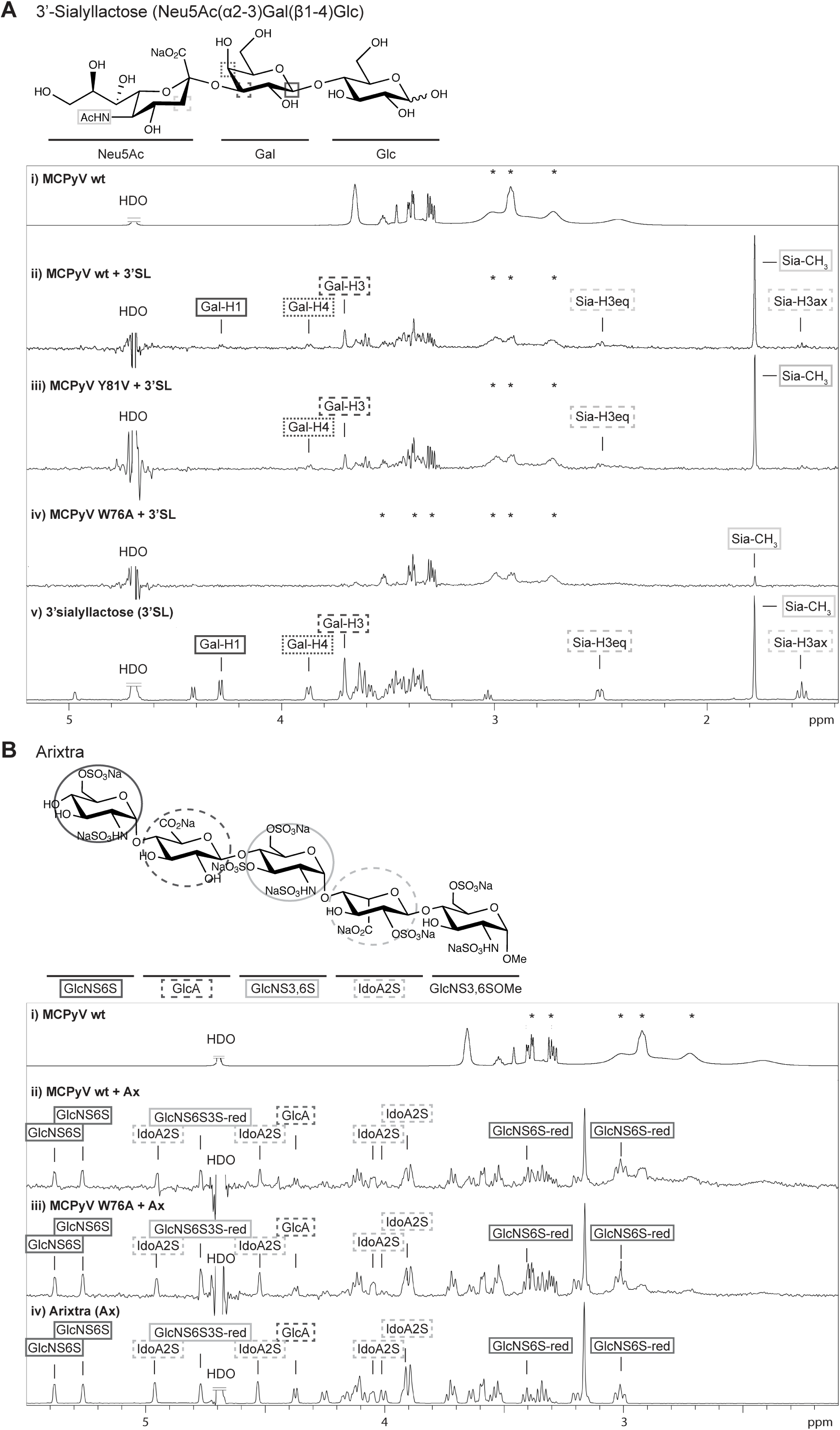
MCPyV sialic acid binding mutants are capable of GAG binding. A) STD-NMR difference spectra of MCPyV VLPs with 3’sialyllactose (3’SL) and reference spectra. From top to bottom: i) VLP ^1^ H 1D reference spectrum; ii) STD-NMR spectrum of WT VLPs with 3’SL exhibiting saturation transfer from the capsid to 3’SL; iii) same spectrum for Y81V mutant VLPs; iv) same spectrum with W76A mutant VLPs showing no transfer to 3’SL; v) 3’SL ^1^H 1D spectrum for comparison. Small molecule impurities are observed in the VLP preparation (i and iv), some of which are buffer components (sharp resonances with asterisks), others are unidentified molecules that are likely associated with the capsids (broad resonances with asterisks, the peak broadening suggests slow molecular tumbling, i.e. association with the VLPs). HDO signals were truncated for the sake of visibility. B) STD-NMR difference spectra of MCPyV VLPs with a GAG pentasaccharide (Arixtra, Ax) and reference spectra. From top to bottom: i) VLP ^1^H 1D reference spectrum; ii) STD-NMR spectrum of WT VLPs with pentasaccharide exhibiting saturation transfer from the capsid to the GAG; iii) same spectrum with W76A mutant VLPs; iv) GAG ^1^H 1D spectrum for comparison. Saturation transfer to the GAG is observed for the WT and the W76A mutant that does not bind to 3’SL (Fig. 8 A iv). HDO signals were truncated for the sake of visibility.

EM analyses showed that the W76A and Y81V mutants readily bound to the A549 cells, presumably through interactions with HSPGs (Fig. 8 B ii, C ii; (30)). Interestingly, both mutants were exclusively observed in the second class of invaginations (i.e., invaginations with a >5 nm gap between the plasma membrane and the virion surface (Fig. 8 B ii, C ii). This observation supports the concept that the first class of invaginations are formed through short range contacts between the virion and sialylated glycolipids (which the mutants fail to bind), while the second class of invagination involves interactions with bulkier HSPGs.

## Discussion

MCPyV is the first human PyV clearly linked to the development of a specific cancer (12). Initial studies on the principal mechanism of infection confirmed the requirement of sulfated glycosaminoglycans for attachment and sialylated glycans for infectious uptake into host cells, which may, in fact, reflect a common mechanism for several PyV (28, 30, 31, 88). To extend our knowledge of the mechanism of initial infection, we addressed additional cellular requirements and routes of virus entry. Our evidence from inhibitor and morphological studies indicates that MCPyV infects cells asynchronously via a caveolar/lipid-raft dependent endocytic pathway that is similar to that used by SV40. After internalization, virus particles were routed to endolysosomal compartments and the ER. Our preliminary evidence suggests that entry of MCPyV depends on progression of the cell cycle, potentially for the nuclear entry step.

The initial step of virus entry is binding to target cells. Previous work suggested that the interaction with sulfated GAGs is required for MCPyV binding, whereas sialic acids have a post attachment role (28). We confirmed that heparin, a highly sulfated GAG, was able to compete for cell surface GAGs thus prohibiting binding and infection of A549 cells. In addition, we addressed the role of GAGs and sialic acids concerning their influence on endocytic uptake. Interestingly, we observed two populations of MCPyV particles on the plasma membrane and in endocytic pits differing in their distance to the limiting membrane, possibly representing the different binding receptor species. In principle, these could reflect binding to proteinaceous receptors with a large ectodomain (wide pits) or gangliosides (sialylated lipids, tight pits), as shown for JC virus uptake via sialylated glycoproteins (89) and SV40 uptake by ganglioside interaction (90). Alternatively, the proteinaceous receptor may constitute HSPGs.

To further unravel the individual roles of sialylated glycans and GAGs during MCPyV entry, two sialic acid binding site mutants (W76A and Y81V, (30)) were subjected to a STD-NMR-based binding assay. These studies confirmed that binding to sialylated glycans was abolished for W76A, but not for Y81V, whereas either mutant retained binding of the HS pentasaccharide. This finding indicates that the binding sites for sialic acid and sulfated GAGs do not overlap and are independent of each other. The most striking difference between wild-type MCPyV and the sialic acid binding site mutants in the TEM experiments was that the mutants were exclusively detected in endocytic pits with the large distances between viral particles and plasma membrane layers. The most likely interpretation of these observations is thus that the large distance reflects binding to a proteinaceous receptor containing sulfated GAGs such as HSPGs, whereas tightly fitted binding to the plasma membrane reflects ganglioside engagement. Since uptake of HSPGs has been suggested to occur primarily through flotillin- and dynamin-dependent endocytosis (91, 92), MCPyV internalization in wider pits via HSPGs seems distinct from the caveolar/lipid raft-mediated internalization required for infection. It thus appears that MCPyV internalization via HSPGs is a dead end. While W76A and Y81V MCPyV were readily observable in endosomes, they were not detected in the ER. Hence, the W76A and Y81V mutations likely rendered the particles non-infectious, as they were unable to engage sialic acid containing gangliosides efficiently and thus were not taken up by an infectious pathway that routes the virus to ER.

Since the sialic acid binding mutants were found in endosomes but not the ER, the interaction with sialic acid receptors is crucial for routing the virus to the ER. This is in line with previous work indicating that branched sialylated glycosphingolipids targets murine polyomavirus (mPyV) and SV40 from endolysosomes to the ER (36, 48, 80, 93). The role of HSPGs remains less clear. It may be that MCPyV interacts with sulfated GAGs simply, because they may serve as an initial high affinity attachment factor, which facilitates later interaction with low affinity glycosphingolipids. An alternative mechanism may be the induction of a conformational change in the virus capsid upon GAG binding that may facilitate uncoating and transfer to the secondary receptor, as it has been described for HPV16, a virus with a similar tissue tropism (51, 94). However, such GAG-induced conformational changes have no precedent among other PyVs. Of note, JC PyV can utilize two distinct pathways for infectious internalization, where one depends on interaction with sulfated GAGs and the other relies on sialic acid binding, this effect could depend on the host cell type (88). As the niche(s) for MCPyV infection remain only partially understood, the role of HSPGs in MCPyV entry may thus depend on specific cell types that are being infected *in vivo.*

Receptor engagement leads to the endocytosis of MCPyV. In line with our initial hypothesis that infectious entry of MCPyV may follow a path similar to that of SV40, our inhibitory experiments showed that infectious entry was independent of clathrin and the activity Na^+^/H^+^-exchanger but required dynamin and cholesterol, as well as actin dynamics. Since MCPyV particles were not found in coated pits or membrane protrusions, entry does not involve CME or macropinocytosis. The combined requirements for dynamins, cholesterol and actin dynamics indicated endocytosis by caveolar/lipid raft mediated endocytosis. Dynamins are also involved in phagocytosis and endocytosis of interleukin-2 (IL-2), both of which also require actin and lipid rafts/cholesterol (32). Phagocytosis occurs only in specialized cells and it is characterized by long outward protrusions that close around large cargos (95). Thus, an involvement of phagocytosis can be excluded similar to macropinocytosis. IL-2 endocytosis, on the other hand, occurs into small, inward budding vesicles, which are mostly formed at the base of an outward protrusion (96). Such initial pits containing MCPyV can be found in the electron micrographs, however in the majority of cases, MCPyV is taken up into small vesicle from flat membrane regions, which makes IL-2- like endocytosis seem unlikely.

Caveolar/lipid raft-mediated endocytosis routes cargos into endosomal compartments (48, 97). We found MCPyV particles in high abundance in endosomal compartments resembling endosomes, lysosomes, multivesicular and lamellar bodies. We also observed small numbers of MCPyV virions in the ER. Retrograde trafficking from endosomes to the ER is a general mechanism employed by PyVs. The low frequency of ER-localized MCPyV virions is similar to prior observations of mPyV entry (93) and hints at a bottleneck for trafficking from endosomes to the ER. Murine PyV as well as JC and BK PyV have been shown to require acidic pH environments in the endolysosomal compartment presumably for a conformational change facilitating membrane penetration after translocation into the ER by a yet unknown mechanism (86, 98, 99). For MCPyV, acidification in the endolysosomal system may serve a similar role.

MCPyV infection is sensitive to DTT treatment (Fig. 1 C) and may therefore be dependent on the redox environment within the ER. In analogy to SV40, uncoating and membrane translocation to the cytoplasm may occur in the ER with the help of the ERAD machinery and cytoplasmic chaperones (39, 40).

As a final step, the viral genome must be delivered to the nucleus, which is the site of early gene expression and replication for PyV. Interestingly, we found that MCPyV infection was sensitive to cell cycle block in S-phase, which indicates that mitosis is required for completion of MCPyV entry. Previous work on HPV16 identified its dependence on mitotic activity of the target cells, which allows delivery of the viral genome to the nucleus upon nuclear envelope breakdown (77, 100). However, SV40 has been described to enter the nucleus of interphase cells through the nuclear pore complex by making use of nuclear localisation signals in the viral capsid proteins after ERAD-dependent partial disassembly (40, 101-103). It remains thus unclear, why MCPyV target cells actively progressing through the cell cycle. Nevertheless, our findings are in line with the importance of WNT signalling during infection of dermal fibroblasts (25).

To our surprise, we regularly observed enveloped particles in endosomal compartments. It remains unclear when and how these virions acquire a membraneous envelope, whether this reflects an antiviral mechanism or whether it is part of the infectious entry mechanisms. It is conceivable that MCPyV uses the ESCRT machinery that generates intraluminal vesicles (ILVs) in multivesicular bodies (104). Alternatively, it may wrap itself in a membrane within the late endosomal/lysosomal compartment. The membrane could theoretically shield the virus from hydrolases, and may give it the time to persist until infection can be completed e.g. by progression through the cell cycle. Alternatively, these enveloped virus particles might arise through an unknown antiviral mechanism. Future studies will address the relevance of such enveloped particles for the infectious route.

In the light of our experiments and previous reports (28, 30), we propose that MCPyV attachment and internalization, at least in the model A549 cell line, is mediated by HS-type GAGs but that additional sialic acid binding is essential to route the internalized virus to the productive infection pathway of retrograde ER trafficking. However, entry into the ER seems to be a bottleneck for infections.

## Materials and Methods

### Cell lines, antibodies, and reagents

HeLa cells were from ATCC. A549 cells were a kind gift from C. Buck (NIH, National Cancer Institute, Bethesda, MD, USA). CV1 cells were a kind gift from J. Kartenbeck (DKFZ, Heidelberg, Germany). BHK Helsinki cells were a kind gift from A. Helenius (ETH Zürich, Switzerland). 293TT cells were a kind gift from J. Schiller (NIH, National Cancer Institute, Bethesda, MD, USA). Aphidicolin, EIPA, cytochalasin D, nocodazole, nystatin, NH_4_Cl, NaClO_3_, heparin, and sodium orthovanadate were from Sigma-Aldrich. Bafilomycin A1, genistein, progesterone were from Applichem. Brefeldin A, cyclosporine A, jasplakinolide, okadaic acid were from Calbiochem. Dynasore was from Merck. Pitstop2 was from Abcam. RedDot2 was from VWR.

### Viruses

MCPyV pseudoviruses (PsV) containing a GFP reporter plasmid (MCPyV-GFP) were produced by transfection of 293TT cells with pwM2m, ph2m and phGluc as described previously (105, 106). In brief, 293TT cells were transfected with the indicated plasmids. After 48 h, cells were harvested and lysed. For optimal maturation, lysates were incubated for further 24 h with 25 mM ammonium sulfate (pH 9.0) (107). MCPyV PsVs were purified on a 25%-39% linear OptiPrep gradient (Sigma-Aldrich). HPV16 pseudoviruses containing a GFP reporter plasmid (HPV16-GFP) were produced by transfection of 293TT cells with p16SheLL and pCIneo as described previously (105, 106). The procedure was similar to MCPyV production above. SV40 and SFV were produced as described previously (40, 108).

### Infection assays upon inhibitor treatment

Infection assays with MCPyV, HPV16, and SV40 were conducted in 96well plates, where 3000 or 10000 A549 cells were seeded in RPMI (5% FCS and 2 mM glutamine) at least 6 h prior to infection. Cells were treated with 80 µL of the following inhibitors at the indicated concentrations diluted in RPMI (5% FCS, 2 mM glutamine) or SV40 infection medium (RPMI with 3% BSA, 10 mM HEPES, pH 6.8): overnight preincubation - nystatin/progesterone, sodium chlorate, and toxin B; 30 min preincubation – bafilomycin A1, brefeldin A, cytochalasin D, dithiotreitol, dynasore, EIPA, genistein, jasplakinolide, NH_4_Cl, nocodazole, okadaic acid, pitstop2, sodium orthovanadate, wortmannin. Virus was diluted in RPMI (5% FCS, 2 mM glutamine) or SV40 infection medium and 20 µL of the inoculum were added to each well. MCPyV samples were incubated for 30 h until the inhibitor dilutions were exchanged to RPMI containing 10 mM NH4Cl and 10 mM HEPES, pH 7.4 and incubated for further 42 h. Samples were fixed by addition of a final 4% paraformaldehyde (PFA) to the wells. Nuclei were stained with RedDot2. Infection scored by analysis of GFP expression by microscopy (Zeiss Axio Observer Z1 equipped with a Yokogawa CSU22 spinning disc module and a CoolSnap HQ camera; Visitron Systems GmbH). Infection with HPV16 was similar to MCPyV but for the exchange of medium to DMEM containing 10 mM NH_4_Cl and 10 mM HEPES, pH 7.4 occurred after 12 h post infection and fixation was done after total 48 h. SV40 inoculum was replaced by fresh DMEM containing 5 mM DTT after 10 h incubation until fixation after 24 h. Cells were incubated with SFV for 4 h until fixation. Samples were fixed as above. For detection of infection, SV40 and SFV samples were stained with an anti-LTag antibody (*sc-20800*) or SFV glycoprotein antibody (110), respectively, and an AF488-coupled secondary antibody; subsequently nuclei were stained with RedDot2. Infection was scored by microscopy (Zeiss Axio Observer Z1 equipped with a Yokogawa CSU22 spinning disc module and a CoolSnap HQ camera; Visitron Systems GmbH). Cell numbers and infection indices were determined by using the MatLab script Infection Counter as described before (40, 48).

Alternatively, the effects of single inhibitors on MCPyV and control virus infection were tested in a flow cytometry-based assay. Here, 3×10^4^ A549 cells were seeded in 24-well plates or 5×10^4^ HeLa cells in 12-well plates or 2.5×10^5^ BHK Helsinki cells in 12-well plates 6-8 h prior to treatment with aphidicolin over night. The compound was renewed before cells were infected with MCPyV, HPV or SFV, respectively. Aphidicolin was renewed daily until fixation of the cell at 72 h p.i. by trypsinization and subsequent addition of 4% paraformaldehyde. The percentage of GFP-positive cells was measured by flow cytometry using a BD FACScalibur.

### Electron microscopy

For negative staining EM of virus particles, about 8×10^6^ PsV in PBS/0.8M NaCl were absorbed for 5 min on formvar coated, carbon sputtered grids. Particles were contrasted for 7 min with 1% phosphotungstic acid. Samples were analyzed directly after drying. The sample was analyzed at 80 kV on a FEI-Tecnai 12 electron microscope (FEI, Eindhoven, Netherlands). Images of selected areas were documented with Olympus Veleta 4k CCD camera.

For ultrathin sectioning transmission EM, virus particles were added for 2 h, 8 h and 16 h to A549 cells before fixation with 2.5% glutaraldehyde in PBS. Samples were post-fixed with 0.5% OsO_4_, block stained with 0.5% uranyl acetate and after dehydration embedded in Epoxyresin. 60nm ultrathin sections were cut and counterstained with uranyl acetat and lead. Images of selected areas were documented with Olympus Veleta 4k CCD camera.

### NMR spectroscopy

NMR experiments were conducted at 283 K using a 600 MHz Bruker Avance spectrometer equipped with a TXI triple resonance room temperature probe head. 3 mm I.D. NMR tubes with 200 µL sample volume were used. Each sample contained 55.6 nM MCPyV VLPs (i.e. 20 µM of the major capsid protein VP1) - either wild-type or mutant - and 1 mM oligosaccharide (either 3’SL or the Arixtra GAG-pentasaccharide). Prior to NMR sample preparation, VLPs were dialysed in Slide-A-Lyzer MINI dialysis devices (Thermo Fisher Scientific) against 150 mM NaCl, 1 mM CaCl_2_, pH 6.0 in D_2_O. 3’SL (Carbosynth) was added from a 40 mM stock solution prepared in pure D_2_O and Arixtra was added from a 7.2 mM stock solution in 150 mM NaCl, 1 mM CaCl_2_, pH 6.0 in D_2_O, dialysed from ready-to-inject syringes (Aspen). For 3’SL-containing samples, off- and on-resonance frequencies in STD-NMR experiments were set to −30 ppm and 7.3 ppm, respectively, while -30 ppm and -0.5 ppm were used for Arixtra-containing samples, owing to the different chemical shift ranges of both glycans. The irradiation power of the selective pulses was set to 57 Hz. The saturation time was 2 s and a total relaxation delay of 3 s was used. A 50-ms continuous-wave spin-lock pulse with a strength of 3.2 kHz was employed to suppress residual protein signals. A total number of 512 scans and a total number of 10,000 points were collected and spectra were multiplied with a Gaussian window function prior to Fourier transformation.

### MCPyV VLP preparation for NMR spectroscopy

MCPyV VLPs were produced essentially according to a published protocol (111). The pwM and ph2m vectors coding for MCPyV VP1 and VP2, respectively, ((27), see also www.addgene.org) were used at a 5:1 ratio for transfection of 293 TT cells. OptiPrep was omitted during the purification and replaced with two CsCl density gradient centrifugation steps.

## Acknowledgements

We would like to thank N. Cordes and E. Weghake (Cellular Virology, Münster, Germany) for technical support during virus production and infection experiments. Thanks also to members of the Schelhaas laboratory for helpful comments on this manuscript. This work was supported by funding to MS by the German Research Foundation (DFG EXC 1003 (partly)) and within the InfectERA initiative by funding from the Federal Ministry for Education and Research (BMBF, 031L0095A). Further support from the DFG to BSB is also acknowledged (1294/3-1 belonging to FOR2328 ‘Virocarb’).

